# Oncogenic RABL6A promotes NF1-associated MPNST progression in vivo

**DOI:** 10.1101/2021.07.07.451475

**Authors:** Jordan L Kohlmeyer, Courtney A Kaemmer, Joshua J Lingo, Mariah R Leidinger, David K Meyerholz, Munir R Tanas, Rebecca D Dodd, Dawn E Quelle

## Abstract

**Background:** Malignant peripheral nerve sheath tumors (MPNSTs) are aggressive sarcomas that display complex molecular and genetic alterations. Powerful tumor suppressors *CDKN2A* and *TP53* are commonly disrupted in these lesions along with *NF1*, a gene that encodes a negative regulator of Ras. Many additional factors have been implicated in MPNST pathogenesis. A greater understanding of critical drivers of the disease is needed to guide more informed targeted therapies for patients. RABL6A is a newly identified driver of MPNST cell survival and proliferation whose *in vivo* role in the disease is unknown.

**Methods:** Using CRISPR-Cas9 targeting of *Nf1*+*Cdkn2a* or *Nf1*+*Tp53* in the sciatic nerve to form *de novo* MPNSTs, we investigated the biological significance of RABL6A in MPNST development. Molecular evaluation of terminal tumors (western blot, qRT-PCR, immunohistochemistry) yielded several insights.

**Results:** Mice lacking *Rabl6* displayed slower tumor growth and extended survival relative to wildtype animals in both genetic contexts. YAP oncogenic activity was selectively downregulated in RABL6A-null, *Nf1*+*Cdkn2a* lesions but not in RABL6A-null, *Nf1*+*Tp53* tumors. Regardless of genetic context, loss of RABL6A caused upregulation of the CDK inhibitor, p27 in tumors. Paradoxically, both models displayed elevated Myc protein expression and Ki67 staining in terminal tumors lacking RABL6A.

**Conclusions:** These findings demonstrate RABL6A is required for optimal tumor progression of NF1 mutant MPNSTs *in vivo* in both *Cdkn2a* and *p53* inactivated settings. However, sustained RABL6A loss may provide selective pressure for molecular alterations, such as Myc upregulation, that ultimately promote an unwanted, hyper-proliferative tumor phenotype akin to drug resistant lesions.

**Importance of the Study:** MPNSTs are aggressive, deadly, and challenging to treat tumors due to location around nerves and high mutational burden. Many factors implicated in MPNST genesis have yet to be fully tested for biological significance in disease formation. We establish a critical physiological role for a new oncoprotein, RABL6A, in promoting NF1-associated MPNST progression. We identify novel RABL6A-regulated pathways that likely contribute to tumor growth, specifically YAP and Myc signaling, and found that sustained RABL6A loss eventually yields more proliferative tumors. We liken RABL6A deficient tumors to those being treated with therapies targeting RABL6A effectors, such as CDKs. Therefore, those lesions should provide a powerful platform to uncover key mediators of drug resistance. Our data suggest oncogenic YAP and Myc could be such mediators of resistance. This study provides a novel system to examine one of the most pressing clinical challenges, drug resistant tumor growth and relapse, in cancer therapy.

## Introduction

Malignant peripheral nerve sheath tumors (MPNSTs) are deadly soft tissue sarcomas that arise from the Schwann cells surrounding peripheral nerves. MPNSTs occur sporadically or in association with the tumor prone, neurological disorder Neurofibromatosis Type I (NF1). Disruption of the *NF1* gene, which encodes neurofibromin, a negative regulator of the Ras oncoprotein, is a key underlying feature of MPNST pathogenesis [1]. In NF1 disease, loss of neurofibromin induces benign neurofibromas (NFs) that can undergo malignant transformation following loss of the *INK4A/ARF* (also called *CDKN2A*) and/or *TP53* tumor suppressors [2]. A number of other factors have been implicated in MPNST development; however, many of them still require biological testing [3]. Thus, while our understanding of MPNST development is continually expanding, much more remains to be learned about the critical alterations driving disease progression. Elucidating these pathways will guide the rational development of targeted therapies to combat MPNSTs, which currently lack effective treatments.

To uncover meaningful alterations driving MPNSTs, genetically engineered mouse models (GEMMs) have been generated to delineate genes and pathways contributing to MPNST biology. Some of the earliest GEMMs to successfully develop MPNSTs involved combined loss of *Nf1* with *Trp53* [4, 5], *Pten* [6] or *Cdkn2a* [7, 8]. Others employed *Sleeping Beauty* transposon-based mutagenesis in p53 mutant mice with overexpressed EGFR to identify cooperating mutations in MPNST pathogenesis [9]. Alterations in Wnt/β-catenin, PI3K-AKT-mTOR, and growth factor receptor signaling pathways were found to promote tumorigenesis in concert with p53 loss and EGFR activation. Most recently, delivery of adenoviruses containing Cas9 and single guide RNAs (sgRNAs) targeting *Nf1* with *p53* into the sciatic nerve of wildtype mice was found to efficiently induce primary MPNSTs [10]. The model produced lesions that are remarkably similar, both genetically and histologically, to human MPNSTs. This provided a unique, rapid and cost-effective tool for investigating MPNST biology. Not only can additional gene targets be concurrently altered by including more sgRNAs in the adenoviral Cas9 construct, but CRISPR editing of *Nf1*+*p53* can be conducted in the peripheral nerves of mice with distinct genotypes.

We recently identified RABL6A, an oncogenic GTPase, as a new regulator of MPNST pathogenesis [11]. RABL6A promotes the proliferation and survival of multiple tumor types, and its increased expression is a marker of poor survival in pancreatic adenocarcinoma [12], breast cancer [13, 14], and non-small cell lung cancer [15, 16]. Histochemical analyses of diverse patient sarcomas, including MPNSTs, showed RABL6A expression is associated with higher tumor grade and shorter time to metastasis [17]. In NF1 patient samples, RABL6A expression is upregulated during the stepwise transformation process with significantly higher levels in MPNSTs versus benign neurofibromas (NFs) [18]. Intermediate lesions, called atypical neurofibromatous neoplasms of uncertain biological potential (ANNUBP), displayed intermediate levels of RABL6A. In MPNST cell lines, RABL6A was found to be necessary for their proliferation and survival, in part through negative regulation of the retinoblastoma (RB1) tumor suppressor pathway. These data suggested a critical role for RABL6A in driving MPNSTs. In this study, we directly examined the biological significance of RABL6A in primary MPNST development and growth using CRISPR-based models of disease lacking either *Nf1*+*p53* or *Nf1*+*Cdkn2a*.

## Materials and Methods

### Primary MPNST generation and growth analysis

Mice were housed in the University of Iowa Animal Care barrier facility in rooms with free access to food and water and a 12hr light-dark cycle. All mouse handling was conducted in strict compliance with the University of Iowa Institutional Animal Care and Use Committee (IACUC) policies under animal care protocol #7112074. These requirements adhere to the National Institutes of Health Guide for the Care and Use of Laboratory Animals and the Public Health Service Policy on the Humane Care and Use of Laboratory Animals. Adenoviruses containing Cas9 and sgRNAs targeting *Nf1*+*Cdkn2a* or *Nf1*+*Trp53* were produced as previously reported [10]. Tumors were initiated by injecting 10μl of CRISPR-Cas9 adenoviruses (4×10^6^ pfu/ul) into the left sciatic nerve of wild-type or *Rabl6* deficient C57BL/6N mice, according to previously established methods [8, 10, 19]. *Rabl6* knockout C57BL/6N mice were generated by the Knockout Mouse Project, as described in [20]. Daily caliper measurements were used to assess tumor growth as previously described [8, 18, 19], which arise as discreet, palpable nodules. Initiation date was determined upon first measurable nodule (125 – 215mm^3^ range), where smallest measurement was 6mm. Tumor volume was calculated using the formula (length x width x thickness x π)/6. Animals were observed for signs of poor health (weight loss, ruffled fur, immobility, and abdominal rigidity). Efforts were made to minimize animal suffering. Mice were euthanized once tumor volume reached 2000mm^3^. Tumors were harvested and split for processing by fixation in 10% neutral buffered formalin or flash frozen in liquid nitrogen.

### Histopathological analysis

Formalin fixed tumors were routinely processed, paraffin embedded, sectioned (~4μm) onto glass slides and hydrated through series of xylene and alcohol baths. 3,3’-diaminobenzidine (DAB, brown staining) was used as the chromogen, and Harris hematoxylin (basophilic staining) was used as the counterstain. Immunostaining for YAP and Ki67 was conducted utilizing validated protocols [17]. Antigen retrieval was performed using citrate buffer (pH 6.0), 110-125°C for 5min, and 20min cool down. Secondary antibodies were obtained from Dako North America, Inc. Slides were reviewed by sarcoma specialist Dr. Munir Tanas.

### RNA isolation and RT-qPCR

RNA was prepared from flash frozen tumors using Qiagen RNAeasy Plus Mini Kit, and cDNA was prepared from 100-200ng RNA using SuperScript III First Strand cDNA Preparation Kit. Diluted cDNA was used for qPCR with gene-specific primers and iQ SYBR Green Supermix reagent in a Bio-Ras CFX96 Real-Time System. qPCR cycling conditions were as follows: (1) denaturation at 95°C for 10min, (2) 40 cycles of 95°C for 15sec and 60°C for 1min. Samples were run in triplicate, and fold change in gene mRNA levels were normalized to *Gapdh* mRNA expression and computed using 2^-ΔΔCt^. Mouse gene-specific RT-qPCR primers for: *Myc* (Fwd: 5’ - CCTGTACCTCGTCCGATTCC – 3’, Rev: 5’ - TTCTTGCTCTTCTTCAGAGTCG - 3’), *Ctgf* (Fwd: 5’ - TTGACAGGCTTGGCGATT - 3’, Rev: 5’ - GTTACCAATGACAATACCTTCTGC - 3’), *Cyr61* (Fwd: 5’ – GTGCAGAGGGTTGAAAAGAAC - 3’, Rev: 5’ - GGAGGTGGAGTTAACGAGAAAC - 3’), and *Gapdh* (Fwd: 5’ - GTTGTCTCCTGCGACTTCA - 3’, Rev: 5’ – GGTGGTCCAGGGTTTCTTA - 3’).

### Western blotting

Flash frozen tumor chunks were lysed in RIPA buffer (50mM Tris, pH 8.0, 150mM NaCl, 1% Triton X-100, 0.1% SDS, 0.5% sodium deoxycholate) containing 1mM NaF, protease and phosphatase inhibitor cocktails (Sigma, P-8340 and P-0044) and 30μM phenylmethylsulfonyl fluoride (PMSF). BCA protein assay was used to determine protein concentrations after lysis. Equal protein amounts were electrophoresed through polyacrylamide gels, and proteins were transferred onto PVDF membranes (Millipore). Membranes were blocked with 5% non-fat milk or 5% BSA in TBST (Tris-buffered saline containing Tween-20) depending on the antibody for 1hr at room temperature. Membranes were incubated with primary antibody solutions overnight at 4°C. HRP-conjugated secondary antibodies and enhanced chemiluminescence (ECL, Amersham, Buckinghamshire, UK) were used to detect proteins. ImageJ (NIH) was used to quantify protein densitometry.

### Antibodies

All antibodies were used in accordance with supplier guidelines. Antibodies used for western blotting include those specific for: c-Myc (1:1000, ab32072) from Abcam, p27 (1:1000, #3686) from Cell Signaling Technology, and p53 (1:200, FL-393, sc-6243), p16 (1:200, F-12, sc-1661), and β-actin (1:500, sc-8432) from Santa Cruz Biotechnology. RABL6A polyclonal antibody (1.5μg/mL) was produced in the Quelle laboratory [12, 21]. Antibodies used for IHC include those specific for: YAP (1:100, sc-15407) from Santa Cruz Biotechnology and Ki67 (1:200, SP6 #16667) from Abcam.

### Statistics

Western data were imaged by scanning densitometry and quantified by ImageJ (NIH). Values for proteins were normalized to expression of the loading control. Differences in levels of protein expression were displayed as fold change relative WT tumors. Quantified data were presented as the mean +/- SD or SEM, as indicated. All *P* values, unless otherwise specified, were obtained by Student’s *t*-test, One-way ANOVA, or Two-way ANOVA and adjusted for multiple comparisons using Dunnett’s test or Bonferroni’s method, as indicated. Overall differences between curves were assessed using generalized linear regressions. An adjusted *P* value less than 0.05 was considered statistically significant.

## Results

### RABL6A promotes tumor growth in two primary MPNST mouse models

In roughly 80% of human MPNSTs, the RB1 pathway is inactivated, primarily by loss of the *CDKN2A* gene locus [1, 2]. RABL6A is a potent oncogene that has been implicated in driving MPNST pathogenesis by negatively regulating the RB1 tumor suppressor pathway [18]. However, the exact contribution of RABL6A expression to MPNST initiation and progression are not known. Therefore, we investigated the consequences of eliminating RABL6A during primary MPNST development. *De novo* MPNSTs were generated via CRISPR-Cas9 targeting of *Nf1*+*Cdkn2a* in wildtype or *Rabl6* knockout C57BL6/N mice (**Figure 1A**). Mice lacking *Rabl6* displayed significantly slower kinetics of tumor growth (**Figure 1B-C**), although no change in tumor initiation rates was observed. Notably, the decreased tumor progression in *Rabl6* deficient mice was associated with a moderate but reproducible extension in survival (median 26.2 days) relative to wildtype controls (median 19.6 days) (**Figure 1D**).

**Figure 1:**
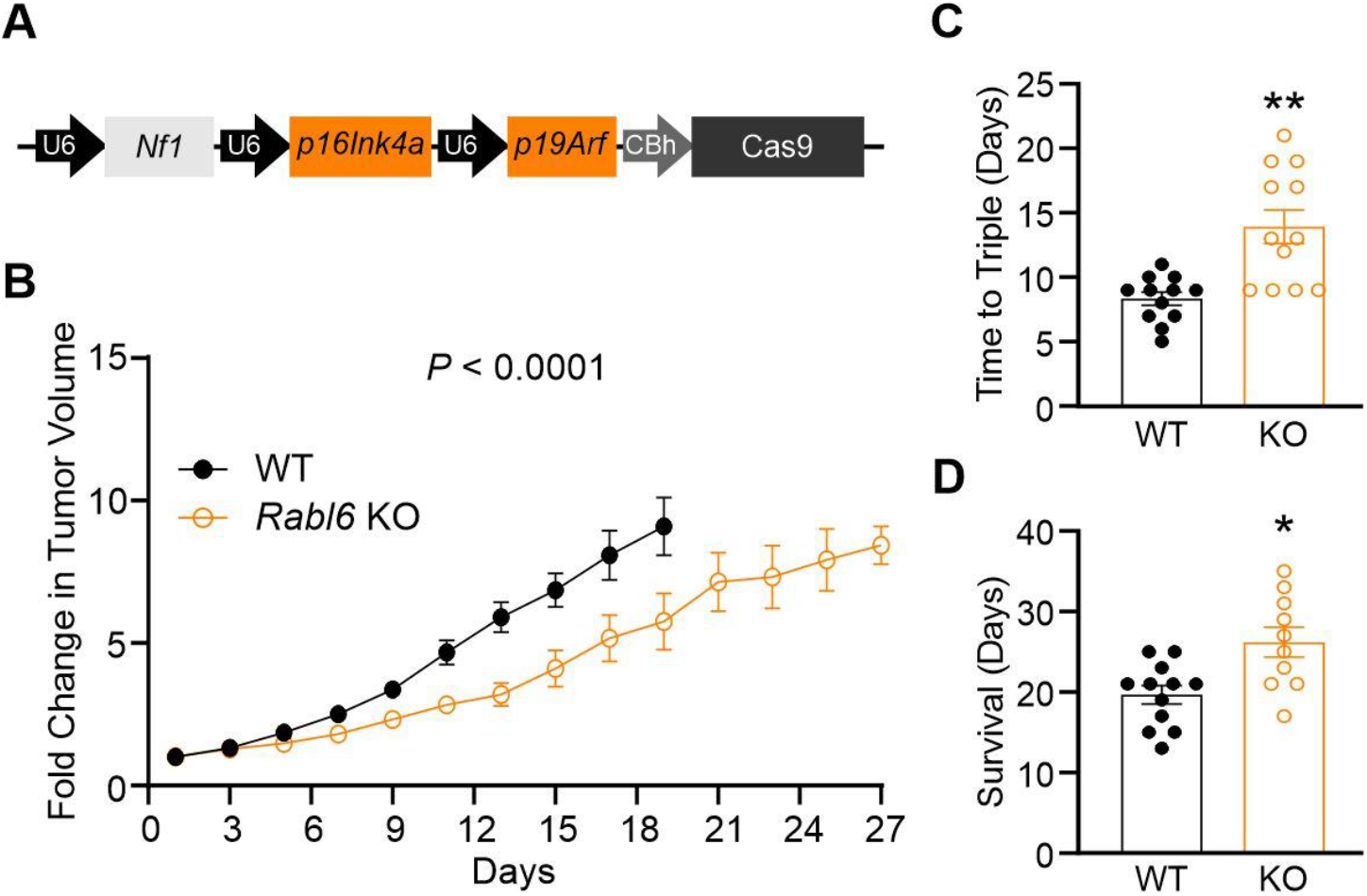
RABL6A promotes MPNST growth in Nf1+Cdkn2a targeted primary MPNST mouse model. Adenoviral packaged CRISPR-Cas9 targeting of *Nf1*+*Cdkn2a* was injected into the sciatic nerve of wildtype or *Rabl6* knockout (KO) C57BL/6N mice to generate primary MPNSTs. (**A**) Schematic of CRISPR-Cas9 targeting construct with sgRNAs against *Nf1* and *p16Ink4a* and *p19Arf* (together comprise *Cdkn2a* locus). (**B**) Fold change in tumor volume shows mice lacking *Rabl6* display slower tumor growth kinetics. (**C**) Time (in days) for tumors to triple in size in WT versus *Rabl6* KO mice. (**D**) Survival (time to maximum 2000mm^3^ tumor volume) of WT versus *Rabl6* KO mice. Error bars, SEM. A: *P* value determined by a generalized linear model to assess the difference between the curves. B: *P* value determined by a generalized linear model to assess the difference between the curves. C-D: *P* value, Student’s *t*-test (*, *P* < 0.05; **, *P* < 0.001).

RABL6A also has functional ties to p53 signaling, which is commonly dysregulated in MPNSTs [2]. RABL6A was first discovered as a new partner of the ARF tumor suppressor [21], which safeguards cells against oncogene activation primarily by activating p53 [22, 23]. Later work showed that RABL6A inhibits p53 activity by directly associating with p53 and Mdm2, enhancing Mdm2-mediated degradation of p53 [24]. Therefore, we examined the biological significance of RABL6A in a second model of *de novo* MPNST that concomitantly targets *Nf1*+*p53* (**Figure 2A**) Tumors were generated and monitored daily in wildtype and *Rabl6* knockout C57BL/6N mice. As observed in the *Nf1*+*Cdkn2a* modified tumors, mice lacking RABL6A displayed significantly reduced tumor progression (**Figure 2B-C**) and extended survival (WT: median 17.9 days, KO: median 23.6 days) (**Figure 2D**).

**Figure 2:**
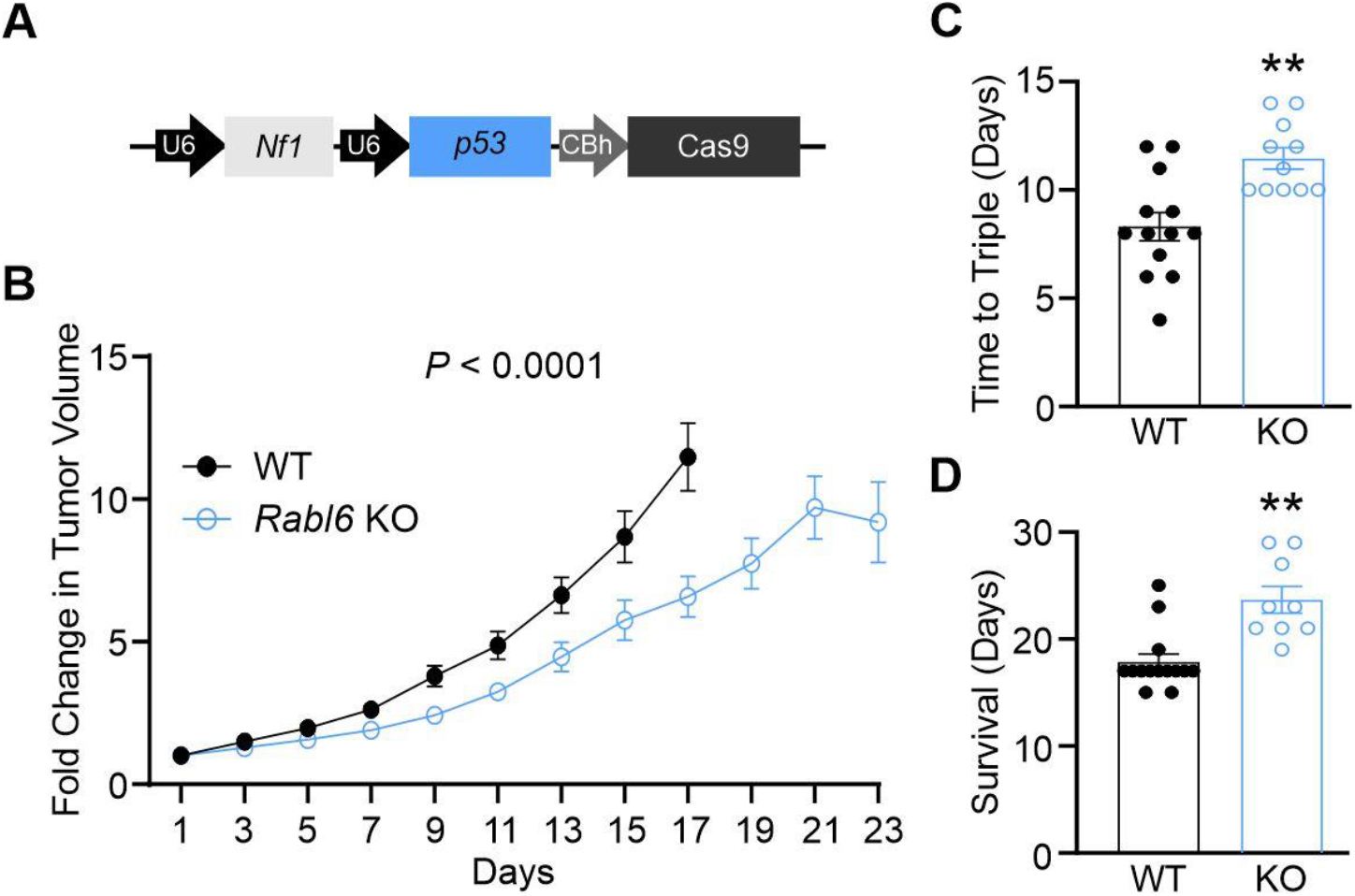
RABL6A promotes MPNST growth in Nf1+Cdkn2a targeted primary MPNST mouse model. Adenoviral packaged CRISPR-Cas9 targeting of *Nf1*+*p53* was injected into the sciatic nerve of wildtype or *Rabl6* knockout (KO) C57BL/6N mice to generate primary MPNSTs. (**A**) Schematic of CRISPR-Cas9 targeting construct with sgRNAs against *Nf1* and *p53*. (**B**) Fold change in tumor volume shows mice lacking *Rabl6* display slower tumor growth kinetics. (**C**) Time (in days) for tumors to triple in size in WT versus *Rabl6* KO mice. (**D**) Survival (time to maximum 2000mm^3^ tumor volume) of WT versus *Rabl6* KO mice. Error bars, SEM. A: *P* value determined by a generalized linear model to assess the difference between the curves. B: *P* value determined by a generalized linear model to assess the difference between the curves. C-D: *P* value, Student’s *t*-test (*, *P* < 0.05; **, *P* < 0.001).

### *Rabl6*-deficient tumors display reduced YAP activity in a context dependent manner

Recent work identified YAP (Yes-Associated Protein), a potent oncogene and transcriptional regulator involved in Hippo signaling, as a driver of MPNST pathogenesis [25, 26]. We identified a significant positive correlation between expression of YAP and RABL6A across numerous types of sarcoma, including MPNSTs [17]. We speculated that RABL6A loss may be associated with reduced YAP expression and activity in our MPNST tumor models. Formalin fixed, paraffin embedded tumors were subjected to IHC for YAP, enabling analysis of its expression and nuclear localization (where it is active). RABL6A depleted *Nf1*+*Cdkn2a* tumors displayed reduced nuclear YAP expression (**Figure 3A**) whereas YAP nuclear levels were unaffected by RABL6A status in *Nf1*+*p53* generated tumors (**Figure 3B**). The reduced nuclear localization of YAP in *Nf1*+*Cdkn2a* tumors lacking RABL6A correlated with diminished mRNA levels of YAP target genes, *Ctgf* and *Cyr61*, compared to tumors from *Nf1*+*Cdkn2a* wildtype mice (**Figure 3C**). Downregulation of *Ctgf* and *Cyr61* was not observed in *Nf1*+*p53 Rabl6* KO tumors, in agreement with immunohistochemical analyses showing no differences between wildtype and *Rabl6* KO tumors. These data suggest a more important role for RABL6A-YAP signaling in *Nf1* mutant, *Cdkn2a*-null lesions compared to *Nf1* mutant MPNSTs bearing p53 alterations.

**Figure 3:**
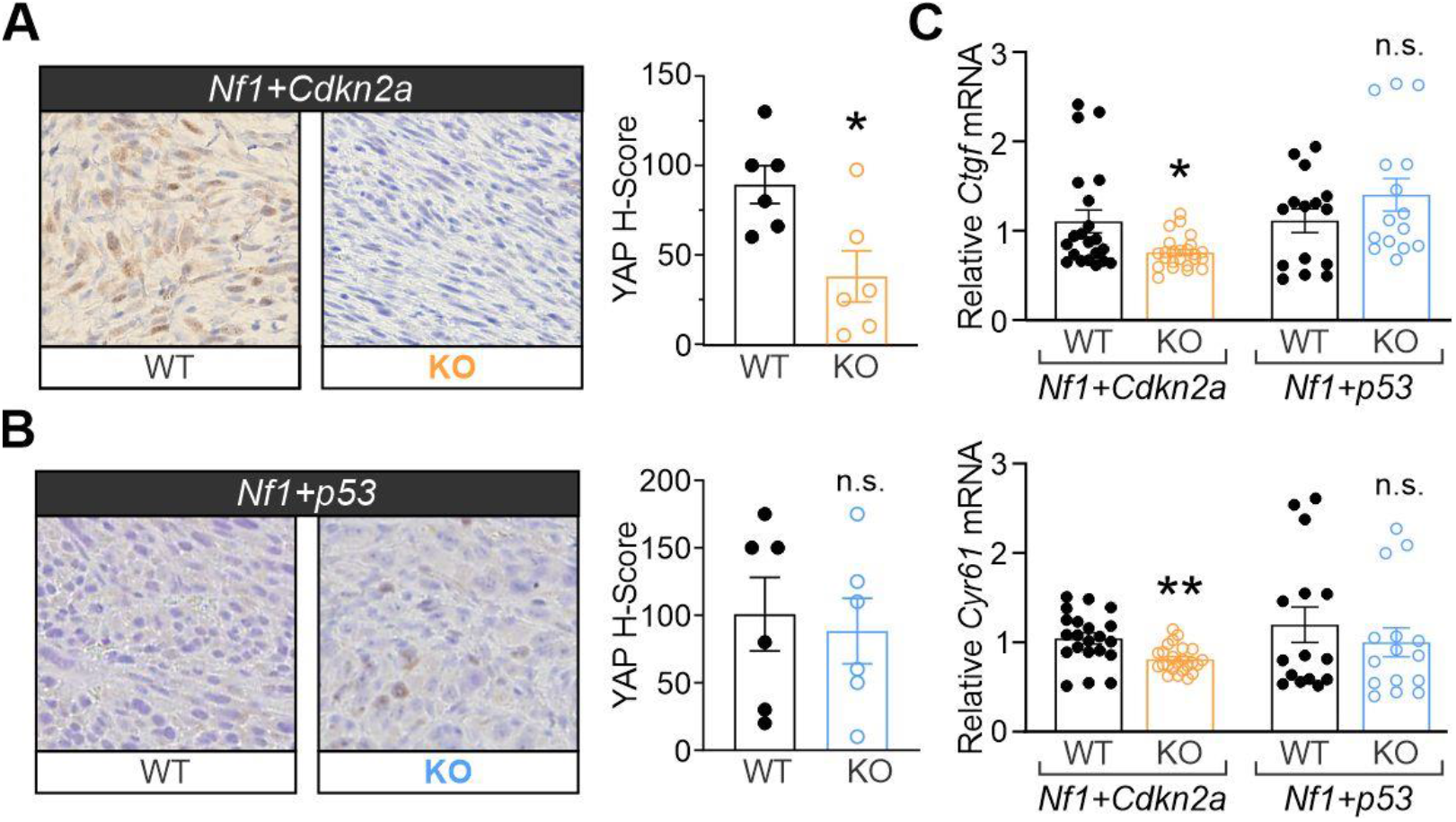
RABL6A-null tumors display reduced YAP activity in a context dependent manner. Representative YAP IHC images (200X magnification) from WT and *Rabl6* KO mice in (**A**) *Nf1*+*Cdkn2a* and (**B**) *Nf1*+*p53* primary MPNSTs with H-Score quantification to the right. (**C**) Relative mRNA levels of YAP target genes, *Ccn2* (Ctgf) (top) and *Ccn1* (Cyr61) (bottom) from WT versus Rabl6 KO Nf1+Cdk2na and Nf1+p53 tumors. YAP expression and activity (measured by downstream target expression) is decreased only in *Nf1*+*Cdkn2a Rabl6* KO tumors compared to WT, whereas *Nf1*+*p53* tumors remain the same. Error bars, SEM. *P* value, Student’s *t*-test (*, *P* < 0.05; **, *P* < 0.01).

### Sustained RABL6A loss leads to paradoxical molecular alterations and restored proliferation

Depletion of RABL6A suppresses *Nf1* mutant MPNST progression in both genetic contexts (**Figures 1-2**). Terminal tumors in each model were harvested and examined for molecular alterations by western blotting and qRT-PCR analyses (**Figure 4**). Western analyses validated loss of p16Ink4a protein in *Nf1*+*Cdkn2a* tumors and p53 protein in *Nf1*+*p53* MPNSTs, as well as RABL6A loss in *Rabl6* KO mice compared to wildtype (**Figure 4A**). Prior work has shown that RABL6A promotes tumor cell proliferation and survival through multiple factors, including p27-CDK-RB1 [18, 27, 28] and PP2A-AKT-mTOR [29] pathways with a role for Myc suggested by evidence that its mRNA and protein expression depend on RABL6A [20, 27]. As recently observed in human MPNSTs [18], *Nf1* mutant mouse MPNSTs lacking RABL6A displayed increased p27 expression compared to wildtype lesions (**Figure 4A-B**). Surprisingly, Myc protein levels were upregulated in *Nf1*+*Cdkn2a* and *Nf1*+*p53* mutant tumors lacking RABL6A (**Figure 4A and 4C**), despite significantly reduced *Myc* mRNA in the same samples (**Figure 4D**). Such data imply the involvement of post-transcriptional events leading to increased Myc protein translation and/or stabilization.

**Figure 4:**
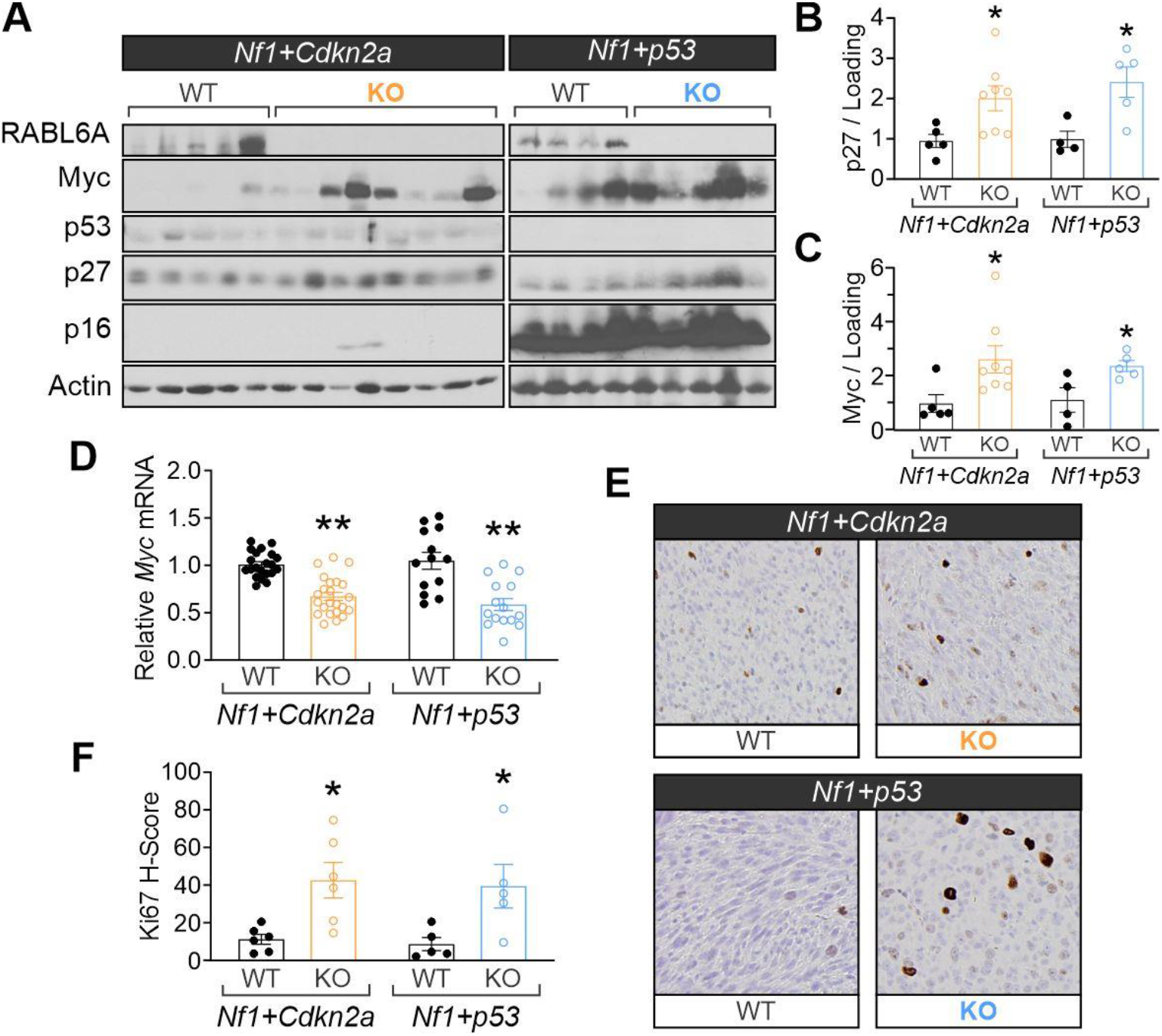
Sustained RABL6A loss leads to paradoxical molecular alterations and restored proliferation. (**A**) Representative western blots confirming knockout of Rabl6, p16, and p53 in respective conditions. *Rabl6* KO mice displayed increased p27 and Myc expression in both *Nf1*+*Cdkn2a* and *Nf1*+*p53* tumors. ImageJ quantification of (**B**) p27 and (**C**) Myc protein expression. (**D**) Significant downregulation of *Myc* mRNA levels in *Rabl6* KO mice. (**E**) Representative Ki67 IHC images (200X magnification) and (**F**) H-scores reveal Rabl6 KO tumors have enhanced proliferation as compared to WT. B-D, F: Error bars, SEM. *P* value, Student’s *t*-test (*, *P* < 0.05; **, *P* < 0.001).

MPNSTs from each model were also examined for potential differences in tumor cell proliferation and survival by immunohistochemical staining. Interestingly, and in agreement with Myc protein upregulation (**Figure 4A-B**), tumors lacking RABL6A displayed higher staining of the proliferation marker, Ki67, relative to wildtype tumors (**Figure 4E-F**). Considered together, these data support a temporal model of how RABL6A loss influences MPNST progression in vivo (**Figure 5**). At first, the absence of RABL6A slows MPNST progression, likely due to upregulation of p27, context-dependent loss of YAP, CDK inhibition and RB1 activation. Over time, however, sustained RABL6A loss provides strong selective pressure for molecular alterations, such as increased Myc signaling, that override the inhibitory effects of RABL6A inactivation and restore tumor cell proliferation (**Figure 5**).

**Figure 5:**
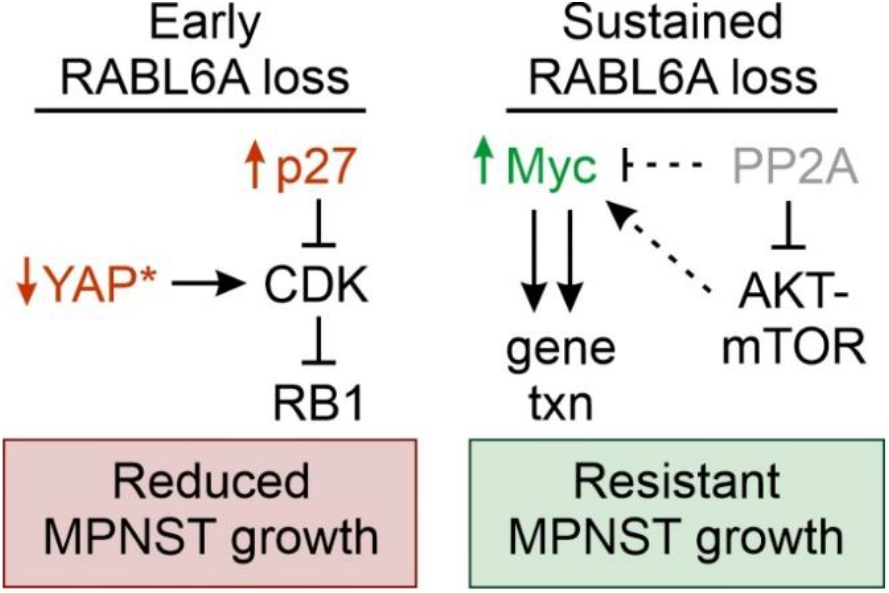
Model of molecular changes in de novo MPNSTs following early versus sustained RABL6A loss. RABL6A-regulated pathways proposed to mediate reduced MPNST growth in the early stages of RABL6A loss (left) versus the highly proliferative, ‘resistant’ tumors that have overcome sustained RABL6A loss (right). Validated changes are highlighted in color. Left, initial loss of RABL6A downregulates YAP expression and signaling (*, selectively in *Nf1-Cdkn2a* inactivated tumors) along with activation of p27-RB1-mediated cell cycle arrest. Right, RABL6A loss is known to increase PP2A tumor suppressor activity, which can destabilize Myc directly or via inactivation of AKT-mTOR. Long-term RABL6A depletion may provide selective pressure for PP2A inactivation (gray and dashed inhibitory bar) and/or AKT-mTOR activation (dashed arrow) that promotes Myc protein stability and the transcription (txn) of its tumor-promoting gene targets.

## Discussion

MPNSTs are difficult to treat cancers due to their location surrounding nerves, aggressive growth and high mutational burden reflecting an extensive number of molecular alterations driving their development. Many factors implicated in MPNST genesis have yet to be fully tested for biological significance in disease formation. Here, we establish a critical physiological role for RABL6A in promoting NF1-associated MPNST progression.

Mounting evidence from earlier work supported the notion that RABL6A would drive MPNST pathogenesis *in vivo*. First, studies of cultured MPNST cells demonstrated RABL6A is an essential regulator of MPNST cell proliferation and survival *in vitro* [18]. RABL6A was found to function, at least in part, by activating oncogenic CDK4/6 through inhibition of p27, thereby disabling RB1-mediated tumor suppression in tumor cells. Those findings, and the fact that p27 loss and RB1 inactivation are hallmark events in MPNSTs associated with worse prognosis [30, 31], predicted RABL6A may be dysregulated in patient tumors. Indeed, RABL6A protein expression is dramatically increased in MPNSTs compared to patient-matched, benign neurofibromas [18]. Notably, RABL6A controls many other cancer pathways besides CDK-RB1, including p53 [24, 27], AKT [29], and Myc [18, 20, 27]. Therefore, we speculated it might have different effects on MPNST growth depending on the genetic context. Using two genetically distinct MPNST models initiated by either *Nf1*+*Cdkn2a* or *Nf1*+*p53* loss, we found that RABL6A is equally required for tumor progression in both settings.

There was one notable difference between the models. YAP nuclear localization and transcriptional activity was significantly reduced in *Nf1*+*Cdkn2a* tumors lacking RABL6A, but not in tumors caused by *Nf1*+*p53* mutation. Mechanisms underlying this difference are not currently known. However, a recent histological analysis of multiple sarcoma subtypes, including MPNSTs, revealed positive correlations between YAP and RABL6A in the patient tumors whereas no direct associations between YAP and p53 were identified [17]. These studies suggest RABL6A may regulate YAP independent of p53 status. Future studies of the RABL6A-YAP link will reveal how biologically relevant this relationship is to MPNST genesis and whether co-targeting YAP with other RABL6A effectors, such as CDK-RB1 signaling, represents a valuable therapy regimen for *Cdkn2a*-altered MPNSTs. This is compelling because YAP activation has been implicated in a number of human cancers [32], most recently including MPNSTs. Wu et al. demonstrated (1) elevated YAP expression in human MPNSTs, (2) YAP hyperactivity in Schwann cells induces MPNSTs, and (3) co-targeting YAP and PDGFR pathways suppressed MPNST growth [25]. More recently, Velez-Reyes et al. employed CRISPR-Cas9 editing of putative MPNST driver genes and identified Hippo-YAP pathway as a likely central driver of MPNST development [26].

RABL6A loss significantly delayed the progression of MPNSTs, but molecular analyses of terminal tumors suggested its sustained absence may induce an unwanted proliferative phenotype similar to the outgrowth of drug-resistant tumors. Specifically, tumors lacking RABL6A exhibited elevated Myc protein levels as well as an increase in the proliferation marker, Ki67. The rise in Myc protein was observed despite reduced *Myc* transcript levels in the same samples, suggesting Myc may be more effectively translated and/or stabilized by post-translational modifications. These data are consistent with the idea that long-term RABL6A depletion, much like anticancer therapies, may provide selective pressure for molecular alterations that override the initial growth inhibitory effects of RABL6A loss. More mouse studies will be needed to test if Myc mediates the hyperproliferative phenotype of *Rabl6*-depleted tumors, but it is likely given the prominent role of Myc in aggressive cancers [33].

The mechanism(s) driving Myc upregulation in tumors with persistent inactivation of RABL6A are not yet known. However, we speculate it may involve dysregulation of the PP2A phosphatase (see **Figure 5**). RABL6A normally inhibits PP2A [29], a prominent tumor suppressor whose dysregulation is seen in many human cancers and is a hallmark of cellular transformation [34, 35]. PP2A directly dephosphorylates Myc and promotes its degradation while also inhibiting AKT and ERK signaling, two downstream effectors of Ras capable of stabilizing Myc [36]. Thus, it stands to reason that initial loss of RABL6A may induce PP2A activity but over time a fraction of cells may escape the growth suppressive effects of PP2A via its inactivation, enabling tumor regrowth. Such alterations would cooperate with hyperactive Ras (due to *Nf1* loss) to promote Myc stability and heighten its transcriptional activity.

Understanding the key alterations that drive tumorigenesis is crucial for developing effective therapies. We recently reported RABL6A negatively regulates the p27-RB1 pathway to promote MPNST pathogenesis, which guided preclinical drug studies that established the efficacy of CDK-targeted therapies in suppressing MPNSTs [18]. Here, we verify the physiological importance of RABL6A signaling in driving MPNST progression in NF1-associated tumors. We identify novel RABL6A-regulated pathways that likely contribute to tumor growth, specifically YAP and Myc signaling, and found that sustained RABL6A loss eventually yields more proliferative tumors. We liken RABL6A deficient tumors to those being treated with therapies targeting RABL6A effectors, such as CDKs. Therefore, those lesions are expected to provide a powerful platform to uncover key mediators of drug resistance. Our data suggest oncogenic YAP and Myc could be mediators of resistance with YAP predicted to play a context-dependent role in *Nf1*+*p53*-mutated tumors. This study provides a novel system to examine one of the most pressing clinical challenges, drug resistant tumor growth and relapse, in cancer therapy.

## Authorship

Conceptualization, JLK and DEQ; Methodology, JLK, CAK, MRL, DKM, MRT, RDD, and DEQ; Formal analysis, JLK, MRT, and DEQ; Writing and Editing, JLK, CAK, DKM, MRT, RDD, and DEQ; Funding acquisition, JLK and DEQ. Study Supervision, DEQ.

## Acknowledgements

We thank Drs. Aloysius Klingelhutz and David Gordon for their constructive thoughts during this study. We are grateful to personnel in the University of Iowa Animal Care facility and Comparative Pathology core facility at the University of Iowa.

